# Intersectional CRISPR-Cas9 genetic targeting reveals acute role of Na_V_1.1 in proprioceptive behavior and function

**DOI:** 10.64898/2026.05.26.727016

**Authors:** Cyrrus M. Espino, Isaac J. Villegas, Serena A. Ortiz, Nadya D. Berber, Julian ANM Halmai, Kyle D. Fink, Theanne N. Griffith

**Affiliations:** Department of Physiology and Membrane Biology, University of California, Davis, Davis, CA, USA; Department of Neurology, University of California, Davis, Davis, CA, USA; Undergraduate Program in Neurobiology, Physiology and Behavior, University of California, Davis, Davis, CA, USA; Howard Hughes Medical Institute

## Abstract

Proprioceptors, a specialized subset of mechanosensory neurons that relay sensory feedback from muscles and tendons, are required for precise, goal-directed movement. Like all neurons, proprioceptors rely on voltage-gated ion channels to generate and transmit electrical signals. Investigating ion channel function in proprioceptors *in vivo* is technically challenging because current approaches to selectively target proprioceptors require the generation of triple transgenic models and creation of both loxP- and Frt-flanked alleles. To facilitate selective targeting of genes within proprioceptors, we employed an intersectional cell-specific gene editing approach that leverages CRISPR/Cas9 and sensory-neuron selective viral capsids. This approach combines single-guide RNA (sgRNA) delivery in sensory neuron-selective adeno-associated viral (AAV) capsids in mice with parvalbumin-driven Cas9 expression. We tested this approach by targeting the voltage-gated sodium (Na_V_) channel Na_V_1.1. Targeting Na_V_1.1 using this approach led to significant motor coordination deficits as early as 3 weeks following sgRNA delivery. Furthermore, whole-cell current clamp recordings from transduced proprioceptors revealed Na_V_1.1 is required for maintaining short-duration action potentials, which would support high-frequency firing typically observed in proprioceptors. Collectively, this study establishes a versatile platform for precise spatiotemporal gene manipulation in otherwise hard-to-access sensory neuron populations, while also providing evidence that Na_V_1.1 is essential for proprioceptor function in adulthood.

**Significance Statement:** Proprioceptors are sensory neurons that relay information about muscle length and force to enable coordinated movement and motor reflexes. Investigating how ion channels contribute to proprioceptor function has been limited by the lack of straightforward and selective genetic tools, which can also confound interpretation of behavioral phenotypes. Here, we developed an intersectional CRISPR/Cas9 strategy that combines sensory-neuron specific viral delivery of sgRNAs with spatially restricted Cas9 expression in mice. Using this method we targeted the voltage-gated sodium channel, Na_V_1.1, which led to persistent motor coordination deficits and impaired proprioceptor action potential waveform, establishing a direct, cell-autonomous role for Na_V_1.1 in proprioceptor function. Thus, our approach provides a flexible platform for spatially and temporally precise gene manipulation in proprioceptors.

## Introduction

The unconscious perception of the body and limbs is known as proprioception. This internal sensation is initiated by a subset of peripheral sensory neurons called proprioceptors, whose cell bodies reside in the dorsal root ganglion (DRG) and mesencephalic trigeminal nucleus of the brain stem (Proske and Gandevia, 2012). Proprioceptors innervate skeletal muscle and tendons to form muscle spindles and Golgi tendon organs that convey mechanical information about muscle velocity and force, respectively (Proske and Gandevia, 2012). Proprioceptive signals are dependent on ion channel activity to transmit information from peripheral sensory end organs to synaptic targets in the central nervous system (Wilkinson, 2022). To study the *in vivo* role of ion channels, transgenic approaches like Cre/LoxP or Flp/FRT systems have been the gold standard for cell type-specific gene manipulations (Park et al., 2011; Maizels, 2013; Yarmolinsky and Hoess, 2015; McLellan et al., 2017). Proprioceptors, however, pose challenges that limit the utility of these systems: (1) The genetic access point for Cre-mediated deletion, Parvalbumin (PV), is also expressed within interneurons of the brain and spinal cord (Ogiwara et al., 2007; Ferguson et al., 2023; Veshchitskii and Merkulyeva, 2023; Zhang et al., 2023); (2) Deletion of ion channels within central PV-interneurons results in motor deficits like those observed in epilepsy (Ogiwara et al., 2007; Meisler et al., 2010; Han et al., 2012; Ferguson et al., 2023); thus any motor phenotypes cannot be solely attributed to proprioceptors; (3) PV is expressed during embryonic development (Veshchitskii and Merkulyeva, 2023); therefore inducible driver lines are needed to investigate these channels in adulthood. One approach to circumvent these limitations would be to create triple transgenic mouse models that require both Cre- and Flp-mediated genetic cassettes (Lehnert et al., 2021). This approach is costly and requires the creation of new mouse lines for each gene of interest. Fortunately, recent advances in molecular genetics and viral delivery systems have laid the groundwork for developing intersectional approaches that are more cost-effective and time efficient.

CRISPR/Cas9 has emerged as an efficient and reliable tool that can generate insertions and deletions (indels) to produce gene loss-of-function (Ran et al., 2013; Wang et al., 2020). This is achieved with single guide RNAs (sgRNAs), which direct the Cas9 endonuclease to induce double-stranded DNA breaks at a specific gene locus (Wang et al., 2020). This gene-editing system has been successfully adapted *in vivo* using different viral approaches. For example, Cre-dependent transgenic mouse lines that enable cell type–specific expression of the Cas9 endonuclease, combined with adeno-associated virus (AAV)-mediated delivery of sgRNAs allow for genome editing in restricted cell populations (Khlghatyan and Beaulieu, 2020). Moreover, single-vector AAV systems incorporating compact Cas9 variants allow for gene editing without the creation of transgenic mouse lines (Hunker and Zweifel, 2020; Hunker et al., 2020; Juarez et al., 2023). While these strategies have been optimized for delivering CRISPR machinery to the central nervous system, targeting peripheral sensory neurons poses additional challenges. Anatomically, DRG neurons are dispersed along the entire rostral-caudal axis, making them less accessible for targeted viral delivery. As a result, systemic administration of non-specific AAVs leads to widespread transduction across the brain, spinal cord, and peripheral tissues, complicating cell-type specific gene targeting. Engineered AAV capsids, such as AAV.PHP.S, have been developed and successfully used to preferentially transduce peripheral sensory neurons *in vivo* (Chan et al., 2017; Asencor et al., 2022).

To overcome the technical challenges associated with targeting proprioceptors, we developed an intersectional CRISPR/Cas9 approach to knockout the voltage-gated sodium (Na_V_) channel, Na_V_1.1, which has been shown to be required for proprioceptive signaling (Espino et al., 2022). We designed and screened sgRNAs against the gene encoding Na_V_1.1, *Scn1a*. Validated sgRNAs were packaged into the AAV.PHP.S vector, and systemically administered to transgenic mice, in which spCas9 is restricted to PV-positive neurons. The intersection of these approaches allows for selective manipulation of *Scn1a* only in proprioceptors. This approach revealed that targeting of *Scn1a* in mature proprioceptors produced motor deficits that were similar, but with notable differences, as those of mice lacking Na_V_1.1 in all sensory neurons (Espino et al., 2022). Whole-cell patch clamp recordings from virally transduced neurons revealed that deletion of *Scn1a* resulted in longer duration action potential, consistent with known roles for Na_V_1.1 in fast-spiking neurons (Kalume et al., 2007; Ogiwara et al., 2007; Han et al., 2012). Collectively, these findings present a tractable intersectional strategy for targeting genes in proprioceptors, which could be adapted to other sensory neuron populations. Furthermore, it demonstrates a requirement for Na_V_1.1 in adulthood for normal proprioceptor function and behavior.

## Materials and Methods

### Experimental model

Both female and male mice were used in this study. All mice used are on a non-congenic C57BL/6J background. Animal use was conducted according to guidelines from the National Institutes of Health’s Guide for the Care and Use of Laboratory Animals and was approved by the Institutional Animal Care and Use Committee of UC Davis (#23049). Parvalbumin^Cre^ (Stock #017320), Rosa26^Cas9^ (Stock #026715) mice were obtained separately from the Jackson Laboratories and bred to generate the Parvalbumin^Cre^;Rosa26^Cas9^ mouse line. Pirt^Cre^ mice were a generous gift from Dr. Xinzhong Dong (Johns Hopkins University; (Kim et al., 2008)). Mice were maintained on a 12hr light/dark cycle, and food and water were provided ad libitum.

### Selection and validation of sgRNAs

Prior to *in vivo* experiments, sgRNAs targeting the mouse *Scn1a* gene were generated using the following *in silico* tools, Synthego ICE design tool (https://design.synthego.com/#/), IDT (https://www.idtdna.com/pages/technology/crispr), and CHOPCHOP (https://chopchop.cbu.uib.no/). Final sgRNAs were selected based on specificity and probability to induce frameshift mutations in *Scn1a.* Each sgRNA was ordered as short oligos (IDT) and subcloned into the pSpCas9(BB)-2A-GFP (PX458) plasmid backbone (Addgene #48138). Subcloned plasmids were transfected into Neuro2A cells using lipofectamine 3000. 2-days post transfection, genomic DNA from cells were isolated (Zymo #D3025) and amplicons surrounding cut site were PCR amplified using an NEB Phusion mastermix kit (#M0531S) with the following primers: *Scn1a* Forward CTCACATGTTTCCTTTGTTGG, Reverse: GCATTCCCCTACACTGTGGT. Amplicons were purified by enzymatic digestion (Qiagen #28104) and sent to Genewiz for sanger sequencing following their standardized sample guidelines with the following sequencing primer: *Scn1a* seq Forward: GTAGTATTTGTGCACATGTTTACCC. Chromatogram files from Genewiz were analyzed using Synthego ICE analysis tool to generate percent insertion and deletion for each sgRNA. SgRNAs with the highest indel percentage were used for downstream experiments.

### Virus production

For behavioral studies, viral particles were prepared in-house as follows: HEK293T cells were triple transfected with the transfer plasmid, and Helper and Rep/Cap (AAV serotype 2/9) plasmids at approximately 80% confluency. Three days post-transfection, cells were harvested and lysed with lysis buffer. Cell pellets and supernatant were processed with PEG8000 (Sigma) for virus precipitation. For purification of AAVs, a 60% iodixanol gradient was used for ultracentrifugation. Fractions of AAV were removed and resuspended in Lactated Ringer’s Solution, then concentrated by centrifugation with 50kDa centrifugal concentrator columns (Sartorius, Fremont, CA, #VS2032). Concentrated AAV viral genomes (vg) were quantified by qPCR using AAVpro Titration Kit (Takara Bio USA, San Jose, CA, #6233). A subset of viral vectors used for *in vivo* validation and *in vitro* electrophysiology were generated by the University of Minnesota Viral Vector and Cloning Core (Minneapolis, MN).

### Retro-orbital injections

Between postnatal day 21 and 25 (P21–P25), mice were injected with either AAV.PHP.S-sgNa_V_1.1 or AAV.PHP.S-sgLacZ for behavioral experiments, or with AAV.PHP.S-sgNa_V_1.1-FLEX-TdTomato and AAV.PHP.S-sgLacZ-FLEX-TdTomato for *in vivo* validation and electrophysiological studies. Injections were performed under anesthesia via the retro-orbital sinus using a 31-gauge, 0.3125-inch insulin syringe. To enhance viral transduction efficiency, mice received two injections (100 µL, 3.8 × 10¹² vg/mouse) administered 24 hours apart.

### Animal behavior

Motor function was tested using three assays: rotarod, open-field, and grip strength. Behavioral testing began prior to viral injections (P21-P25) to obtain a baseline measure of motor performance. Subsequent behavioral testing was carried out 3- and 6-weeks post viral administration. Behavioral assays were conducted from least to most invasive in the order of open field, grip strength, and rotarod. For the open field, mice were acclimated to the behavior room for 1 h prior to testing. The open field apparatus consisted of a white square box with dimensions of 15×15×20 inches. A camera suspended above the open field tracked animal movement for a single 10 min period using ANY-maze software. Following testing in the open field, mice were transported to a separate behavior room and allowed to acclimate for 1 h before being assayed in the grip strength and rotarod tests. A grip strength apparatus (IITC Life sciences, Woodland Hills, CA) with a metal grate was used. Mice held from the tail were placed on the metal grate and pulled horizontally away from apparatus once all four paws touched the grate. Mice were assayed across 6 trials with 5 min intervals between trials. For motor coordination analyses, a rotarod machine (IITC) that has an accelerating rotating cylinder was used. The averages of three trials across three consecutive training days were recorded.

### Tissue processing

At least 3 weeks post injection PV^cre^;Rosa26^Cas9^ mice were anesthetized using a ketamine/xylazine cocktail and transcardially perfused with PBS followed by 4% PFA. Brains were harvested and post fixed in 4% formaldehyde overnight following cryoprotection in a 30% sucrose solution for 2-3 days at 4°C. Spinal cord was harvested in PBS followed by a post fix for 30 minutes then washed in PBS before incubation in 30% sucrose overnight at 4°C. DRG were harvested from all spinal levels and fixed in 4% formaldehyde for 15 min at 4°C and then incubated in 30% sucrose for 2-4 h at 4°C. Following cryoprotection, all tissues were embedded in optimal cutting temperature (Fisher #4585) and stored in −80°C until sectioning.

### Immunohistochemistry

Immunohistochemistry of brain (45μm), spinal cord (30μm), and DRG (25μm) sections was performed using the following primary antibodies: Rabbit anti-DsRed (1:3000, Takara Bio, 632496), guinea pig anti-VGluT1 (1:8000, Zuckerman Institute, 1705), and chicken anti-parvalbumin (1:1000, Zuckerman Institute, 1664) Secondary antibodies used were as follows: anti-rabbit 594 (1:1000, ThermoFisher, A32740), anti-guinea pig 488 (1:1000, ThermoFisher, A11073), and anti-chicken 647 (ThermoFisher, A32733). All specimens were imaged in three dimensions on an Olympus FV3000 confocal microscope. Images were analyzed using ImageJ software.

### DRG culture preparation

DRG were harvested from all spinal levels of adult PV^cre^;Rosa26^Cas9^ mice of both sexes 3 weeks post retro orbital injection and transferred to Ca^2+^-free and Mg^2+^-free PBS solution (Invitrogen, 14170-112). Upon isolation, processes were trimmed, and ganglia were transferred into collagenase (1.5 mg/mL; Type P, Sigma-Aldrich, 11213865001) in HBSS for 20 min at 37°C followed by TrypLE Express (ThermoFisher, 12605-010) for 3 min with gentle rotation. TrypLE was neutralized with 10% horse serum (heat-inactivated; Invitrogen, 26050-070) and supplemented with culture media (MEM with L-glutamine, Phenol Red, without sodium pyruvate, ThermoFisher, 11095-080), containing 10,000 U/mL Penicillin-streptomycin (ThermoFisher, 15140-122), MEM Vitamin Solution (Invitrogen, 11120-052), and B-27 supplement (ThermoFisher, 17504-044). Serum containing media was decanted and cells were triturated using a fire-polished Pasteur pipette in the MEM culture media described above. Cells were resuspended and triturated using a plastic pipette tip. Cells were plated on glass coverslips that had been washed in 2M NaOH for at least 4 h, rinsed with 70% ethanol, UV-sterilized, and treated with laminin (0.05 mg/mL, Sigma-Aldrich, L2020-1MG) for 1 hour prior to plating. Cells were then incubated at 37°C in 5% CO_2_. Cells were used for electrophysiology experiments 14-36 h post-plating.

### *In vitro* electrophysiology

Whole-cell current-clamp recordings were made from dissociated DRG neurons using patch pipettes pulled from Model P-1000 (Sutter Instruments). Patch pipettes had a resistance of 3-5 MΩ when filled with an internal solution containing the following (in mM): 120 KMethylsulfonate, 10 NaCl, 10 KCl, 5 EGTA, 0.5 CaCl_2_, 10 HEPES, and 2.5 MgATP, pH with KOH to 7.2. Seals and whole-cell configuration were obtained in an external solution containing the following (in mM): 145 NaCl, 5 KCl, 10 HEPES, 10 Glucose, 2 CaCl_2_, 2 MgCl_2_, pH 7.3 with NaOH, osmolarity ∼320 mOsm. Upon obtaining whole-cell configuration, neurons were injected with varying amounts of current (−200 to −800 pA) to stabilize resting membrane potential to −60mV prior to experimentation. The calculated liquid junction potential was +0.41 mV and was not accounted for. Current clamp recordings were conducted at 37°C. Bath temperature was controlled and monitored using CL-100 (Warner Instruments).

### Data acquisition and analysis

Voltages were acquired using pClamp software v11.2 (Molecular Devices). Recordings were obtained using an AxoPatch 200b patch-clamp amplifier and a Digidata 1550B and filtered at 5 kHz and digitized at 10 kHz. Action potential threshold was calculated as the membrane potential at which the first derivative of the somatic membrane potential (dV/dT) reached 10 mV ms^-1^ (Griffith et al., 2019; Kress et al., 2008). Current-clamp experiments were analyzed with Clampfit software v11.2 (Molecular Devices).

### Fluorescence-activated cell sorting

On the day of sorting, DRG were isolated from 3 mice per experimental condition and stored in DRG media that contained the following: MEM (Gibco #11095-080), PenStrep (Thermo 15140122), MEM Vitamins (Gibco 11120052), and B27 Supplement (#17504044). Following isolation DRG were transferred to a digestion solutions containing: 5mg/ml Dispase (Gibco 17105-041), 2mg/ml Collegase P (Sigma-Aldrich #11249002001), and 0.1 mg/ml DNAse (Sigma DN25) in HBSS for 20 minutes at 37°C under constant rotation. Following incubation digestion solution was centrifuged at 1000 g for 5 min and cellular pellet was resuspended and triturated in 1 mL of DRG media. Cell suspension was then passed through 70 μm filter that we prerinsed with 1 mL of DRG media. Cell suspension was centrifuged at 1000 g for 5 min and supernatant was discarded. Cells were then resuspended in 500 mL of FACS solution containing: 10mg/ml BSA (Sigma A7906-50G), 1M HEPES, 1 x PenStrep (Thermo 15140122), 0.5 mg/ml DNase in DMEM. Samples were transferred on ice immediately to FACS core facility at UC Davis. A Sony MA 900 cell sorter with a 100μm nozzle was used. DAPI was added right before sorting to exclude dead or damaged cells. Cells were sorted based on TdTomato^+^ and DAPI^-^ signals and were sorted into 1 mL of DRG media. Sorted cells were immediately processed for DNA isolation (Qiagen 80204) and stored at −20°C.

### Whole genome amplification and amplicon sequencing

DNA isolated from sorted cells were subjected to whole genome amplification (WGA) to increase DNA for validation. A New England Biolabs kit (phi29-XT E1604S) was used according to manufacturer protocol. WGA DNA was diluted in nuclease free water and used for the generation of amplicons for amplicon sequencing. For amplicon next-generation sequencing, a ∼200 bp amplicon surrounding the edit was amplified from genomic DNA using PCR primers with five base pair unique molecular indices, or barcodes, attached to the 5’ end of the forward primer. PCR amplicons were purified with the QIAquick PCR Purification Kit (Qiagen, Germantown, MD, USA,#28106) according to the manufacturer protocol. Equal amounts of purified PCR amplicons were multiplexed into a tube with a total volume of 36 µL, with no more than four samples per tube. The Massachusetts General Hospital DNA Sequencing Core performed the library preparation and CRISPR sequencing.

### Experimental design and statistical analysis

Summary data are presented as mean ± SD, from *n* cells, or *N* animals. For quantitative immunolabeling experiments at least 3 biological replicates per condition were used. Statistical differences were determine using parametric tests for normally distributed data and non-parametric tests for data that did not conform to Gaussian distributions or had different variances. Statistical tests are listed in *Results* and/or figure legends. Statistical significance in each case is denoted as follows: *p <0.05, **p < 0.01, ***p < 0.001, and ****p < 0.0001. Statistical tests and curve fits were performed using Prism 11.0 (GraphPad Software).

## Results

### *In vitro* validation of sgRNAs targeting *Scn1a*

To impair Na_V_1.1 function in proprioceptors, we first designed and screened candidate single-guide RNAs (sgRNAs) targeting *Scn1a* of the *Mus musculus* genome. We chose to target Na_V_1.1 based on previous findings indicating *Scn1a* is an abundantly expressed Na_V_ isoform in proprioceptors (Espino et al., 2022). Six candidate sgRNAs were chosen based upon their predicted editing efficiencies and proximity to the 5′ coding region, which maximizes the likelihood of generating loss-of-function protein truncations (Wang et al., 2020)Fig. 1A). Each sgRNA was individually subcloned into a plasmid backbone containing the *Streptococcus pyogenes* Cas9 nuclease (SpCas9) (sgRNA-CMV-SpCas9-GFP). Subcloned constructs were transfected into Neuro2A cells, and bulk genomic DNA was harvested 48 h post-transfection (Fig. 1B). DNA sequences surrounding the predicted cut site for each guide were analyzed by quantifying the prevalence of indels (insertions and deletions) compared to non-transfected control DNA. All candidate guides produced a significant increase in indel frequency relative to non-transfected controls (Fig. 1C). It is likely that the percentage of indels achieved by each sgRNA is an underestimation of the true mutagenic efficiency of each, as DNA from cells that were not transduced by plasmid was also included. Among the screened guides, sgNa_V_1.1_4 (hereafter referred to as sgNa_V_1.1) yielded the highest mean indel rate of 43% (Fig. 1C). Sanger sequencing revealed a diversity of indels that resulted in a premature stop codon 16-19 amino acids downstream of cut site (Extended data Fig. 1-1) Mapping the sgNa_V_1.1 DNA sequence onto protein coding region revealed targeting of exon 1 of the *Scn1a* gene. This region encodes for protein sequences corresponding the N-terminal region of the Na_V_1.1 channel (Fig. 1D); therefore, a premature stop codon induced by Cas9-mediated cleavage at this location would result in a non-functional protein.

**Figure 1.**
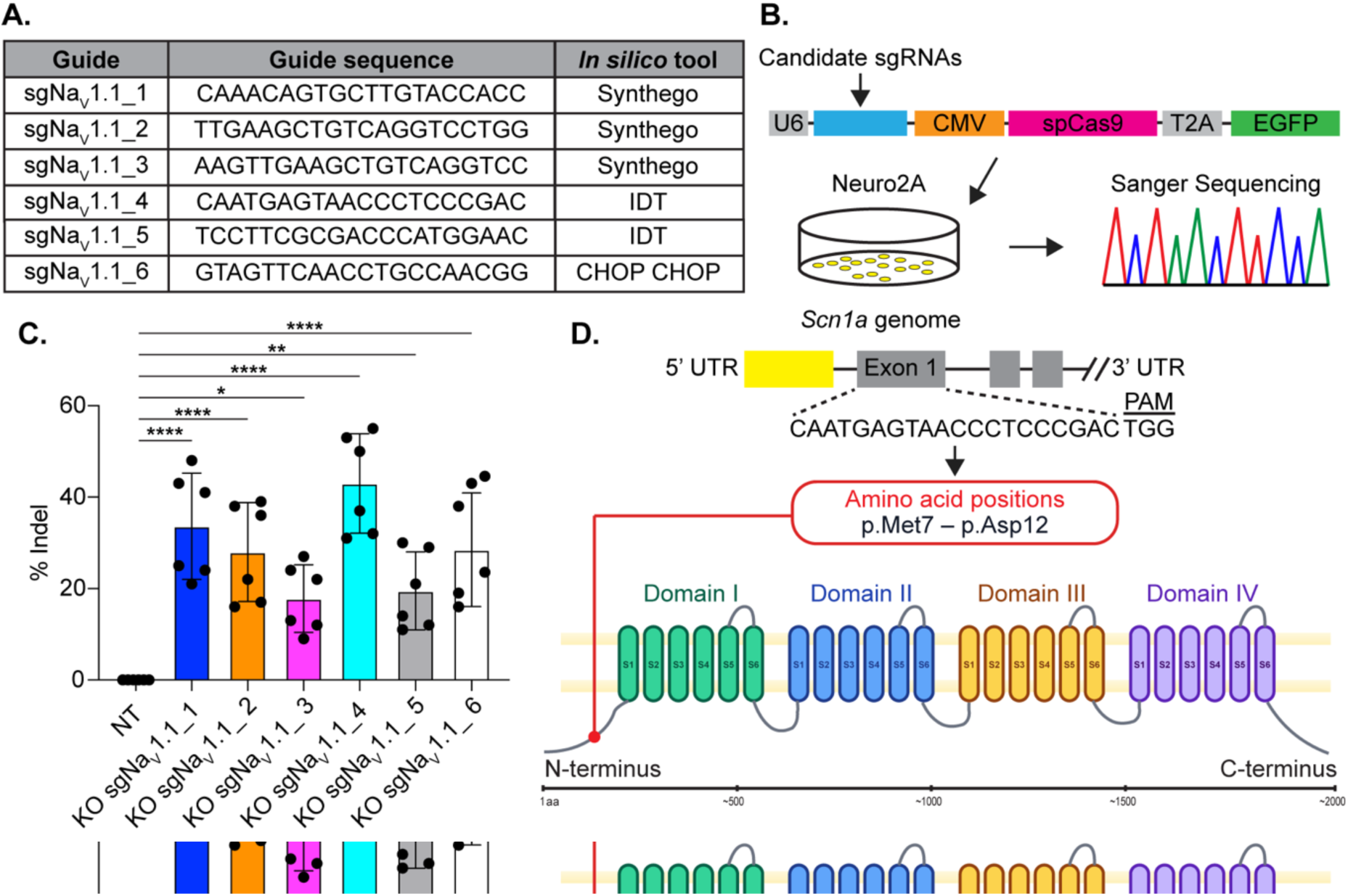
Validation of sgRNAs targeting Na_V_1.1. (A) Table of candidate sgRNA generated using *in silico* tools. (B) Schematic of the experimental workflow for validating sgRNAs in Neuro2A (N2A) cells. Candidate guides were subcloned into a plasmid backbone containing SpCas9. N2A cells were transfected, and 48 hours post-transfection, genomic DNA was isolated and subjected to Sanger sequencing. (C) Quantification of insertions and deletions (indel) frequency from bulk-transfected N2A cells. (D) Cartoon representation of location of sgNa_V_1.1_4 cut site. SgNa_V_1.1_4 targets exon 1 of the *Scn1a* coding region. sgNa_V_1.1_4 targets the N-terminal region in Na_V_1.1 protein (red dot). Statistical significance in (C) was determined using a one-way ANOVA with Dunnett’s post hoc test. Bars denote ± SD with filled circles show data from individual transfections.

### Viral transduction of AAV.PHP.S-FLEX throughout nervous system

We next sought to validate the specificity of the sensory-neuron selective AAV capsid, AAV.PHP.S (Chan et al., 2017), in our experimental paradigm. To do this we generated a transgenic mouse line that expresses spCas9 in all parvalbumin expressing neurons (Parvalbumin^Cre^;Rosa26^Cas9^, PV^Cas9^). We delivered a Cre-dependent fluorescent version of AAV.PHP.S (AAV.PHP.S-FLEX-TdTomato) and examined TdTomato (TdT) expression in DRG, brain, and spinal cord 3 weeks following retro-orbital injection. In DRG sections, we observed TdT expression that colocalized with proprioceptors immunolabeled with anti-PV antibodies (Fig. 2A-C). Quantification of the number of TdT-positive neurons with PV-immunolabeling found on average 71% of PV-positive neurons were transduced with AAV (Fig. 2D). This transduction efficiency is in line with previous reports of this capsid (Chan et al., 2017; Asencor et al., 2022). Additionally, we also examined cerebellar Purkinje neurons, which are also known to express parvalbumin (Brandenburg et al., 2021). We observed sparse labeling of PV-positive Purkinje neurons with TdT (Fig. 2E-G) suggesting viral transduction outside of sensory neurons; however, this labeling was minimal, with an average transduction rate of <1% across quantified cerebellar sections (Fig. 2H). Furthermore, previous studies have shown that PV labels populations of spinal cord neurons (Veshchitskii and Merkulyeva, 2023). TdT labeling appeared as punctate structures, consistent with proprioceptor axonal boutons (Fig. 2K). To confirm this, we co-labeled with antibodies against vesicular glutamate transporter 1 (VgluT1), a marker for proprioceptor synaptic terminals in the ventral spinal cord (Wu et al., 2004) and found that TdT and VgluT1 immunoreactivity were colocalized (Fig. 2I-L). We did not observe any TdT-positive signal that colocalized with PV-expressing neurons, suggesting no off-target transduction in the spinal cord. Lastly, we did not observe any TdT expression in the neocortex, where PV labels populations of cortical interneurons (Ogiwara et al., 2007; Ferguson et al., 2023, Fig. 2M-O). Collectively these data validate AAV.PHP.S for downstream *in vivo* experiments.

**Figure 2.**
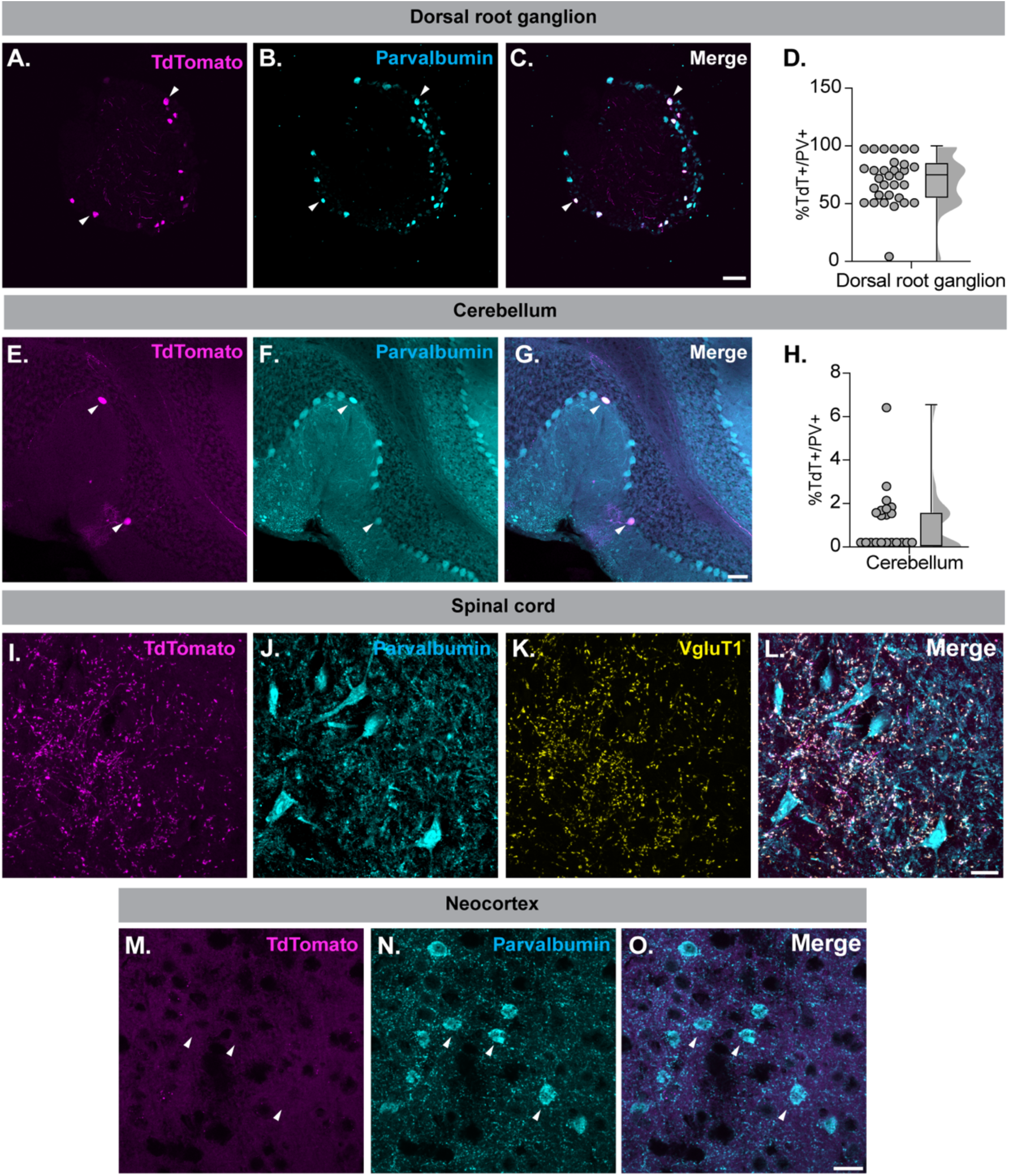
Validation of AAV.PHP.S viral tropism. (A-C) Representative confocal images of a DRG section (25μm) labeled with anti-Dsred (magenta) and anti-parvalbumin (cyan) antibodies. Scale bar set to 100μm. (D) Quantification of the percentage PV-expressing neurons that also express TdTomato. Each dot represents a single DRG section. (E-G) Representative confocal images of cerebellar sections (45μm) labeled with anti-Dsred (magenta) and anti-PV (cyan) antibodies. Scale bar set to 25μm. (H) Quantification of TdT+/PV+ cells in cerebellar sections. Each dot represents a TdT+ neuron across 7 slices (I to L) Representative confocal images of spinal cord sections (30μm) labeled with anti-Dsred (Magenta), anti-PV (cyan), and anti-VgluT1 (yellow) antibodies. Arrows indicate lack of colocalization between PV expressing spinal neurons and TdT. Arrow heads indicated colocalization of TdT and VgluT1 suggesting putative sensory axon terminals. Scale bar set to 25μm. (M to O) Representative confocal images of Neocortical section (45μm) labeled with anti-Dsred (Magenta) and anti-PV (cyan) antibodies. Arrow heads indicate lack of colocalization of PV and TdT in neocortex. Scale bar set to 25μm. At least 3 biological replicates were used for all analyses.

### Confirmation of genomic edits *in vivo*

We next sought to determine whether viral delivery of sgNa_V_1.1 would lead to significant mutagenesis in the *Scn1a* locus, *in vivo*. To do this, we packaged sgNa_V_1.1 and sgLacZ into AAV.PHP.S-FLEX-TdTomato to isolate PV+ cells using fluorescent activated cell sorting (FACS). AAV (3.8×10^12^ vg) was delivered by retro-orbital injection twice 24 h apart. 3 weeks post-injection, DRG were harvested and suspended into single cells. Highly expressing TdT cells were sorted separately, and dead or damaged cells were excluded based on DAPI fluorescence. Following sorting, DNA was extracted and whole-genome amplification (WGA) was performed to enhance the quantity of genomic DNA for downstream processing. This step is consistent with previous studies utilizing similar *in vivo* CRISPR knock out approaches (Hunker and Zweifel, 2020; Hunker et al., 2020; Juarez et al., 2023). DNA sequences flanking the sgNa_V_1.1 cut site were amplified from both sgNa_V_1.1-injected mice, and mice injected with a control sgLacZ viral construct, and subsequently sent for deep sequencing (Fig. 3A). As expected, amplicon sequencing of genomes from mice injected with sgLacZ did not detect the presence of indels (Fig. 3B). Amplicon sequencing of genomes from mice injected with sgNa_V_1.1 revealed 5.42% of amplicons single base pair insertion around the cleavage site (Fig. 3B). Furthermore, we observed double and single base pair substitutions at a rate of 1.4% and 1.13%, respectively. All together the observed indel frequency was 7.95% compared to wildtype amplicons (Fig. 3B). Similar to *in vitro* screen, the indels produced by sgNa_V_1.1 led to a premature stop codon 16-19 amino acids downstream of the cut site (Extended data Fig. 3-1A). It is notable that this indel frequency is much lower than previous studies in the brain, which report a ∼90% indel rate (Hunker et al., 2020; Juarez et al., 2023). We tried an alternative approach, whereby we separately generated an additional transgenic mouse in which Cas9 is expressed in all sensory neurons (Pirt^Cre^;Rosa26^Cas9^, Pirt^Cas9^) (Fig. 3C). Using the same injection and validation pipeline, we observed a similar collection of single and double base pair insertions and substitutions that totaled to an indel frequency of 6.83% compared to reference genomes (Fig. 3C). Similar premature stop codons at 16-19 amino acids were predicted based on indels (Extended data Fig. 3-1B). We predict the source of this low indel frequency is due to the heterogenous nature of the DRG cell suspensions. Indeed, in addition to neurons, DRG suspensions are known to contain significant populations of pericytes, macrophages, satellite glial, Schwann and endothelial cells, totaling nearly 50% of non-neuronal cells (Sharma et al., 2020; Tonello et al., 2023), which are adherent to DRG neurons. These cells, however, would not be transduced by AAV.PHP.S or express Cas9. We therefore suspect that the reported indel frequency is an underestimation of the true rate of mutagenesis *in vivo* and subsequently pursued downstream behavioral and electrophysiological studies.

**Figure 3.**
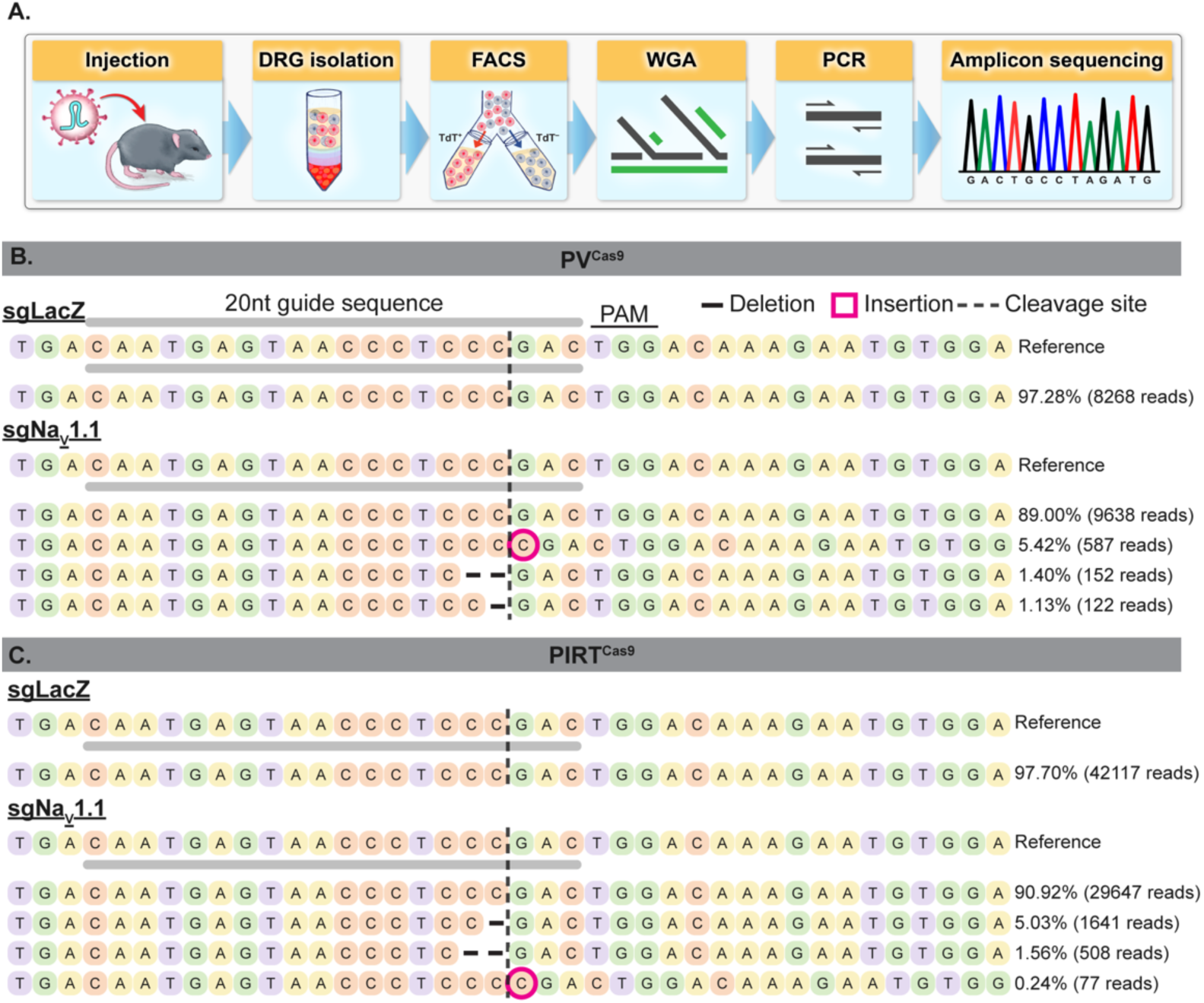
Validation of mutagenesis, *in vivo*. (A) Cartoon schematic depicting experimental workflow to isolate DRG, sort TdT+ cells by FACS, and preparation for amplicon sequencing (B-C) CRISPResso data of top sequenced alleles from amplicon sequencing of the target locus in (B) PVCas9 mice and (C) PirtCas9 mice transduced with AAV.PHP.S-FLEX-TdT containing either sgLacZ controls or sgNaV1.1. Each row shows the 40-nt window centered on the predicted Cas9 Cleavage site (dashed vertical line). The 20-nt guide sequence and PAM site are indicated above the reference sequence. Single-base substitutions relative to reference are highlighted in gray and insertions in magenta. Allele frequencies and absolute read counts (out of total aligned reads per condition) are listed to the right of each row.

### Targeting *Scn1a* in proprioceptors reveals acute role in motor coordination

We next asked whether knockout of Na_V_1.1 in adult proprioceptors would reveal an acute role the channel in proprioceptive driven behaviors. We packaged sgNa_V_1.1 or control sgLacZ into AAV.PHP.S, generating AAV.PHP.S-sgNa_V_1.1 and AAV.PHP.S-sgLacZ (Fig. 4A). Baseline motor performance was assessed at postnatal day (P) 21–25, before viral delivery, and re-evaluated 3- and 6-weeks post-injection (Fig. 4A). Mice receiving sgNa_V_1.1 displayed notable hind-limb motor deficits compared with sgLacZ controls as early as 3 weeks post injection (Fig. 4B-C). These deficits were distinct than those that had been previously reported in mice lacking Na_V_1.1 in all peripheral sensory neurons (Pirt^Cre^;Na_V_1.1^fl/fl^, Espino et al., 2022). We quantified the observed motor deficits in sgNa_V_1.1 or sgLacZ mice on an accelerating rotarod (Fig. 4D). Prior to injection, both groups showed increased latencies to fall from the rotarod, indicative of motor learning and increased coordination with training. 3 weeks post-injection, however, sgNa_V_1.1 mice fell off the rotarod significantly faster than sgLacZ controls, returning to day-1 baseline performance (Fig. 4D).

**Figure 4.**
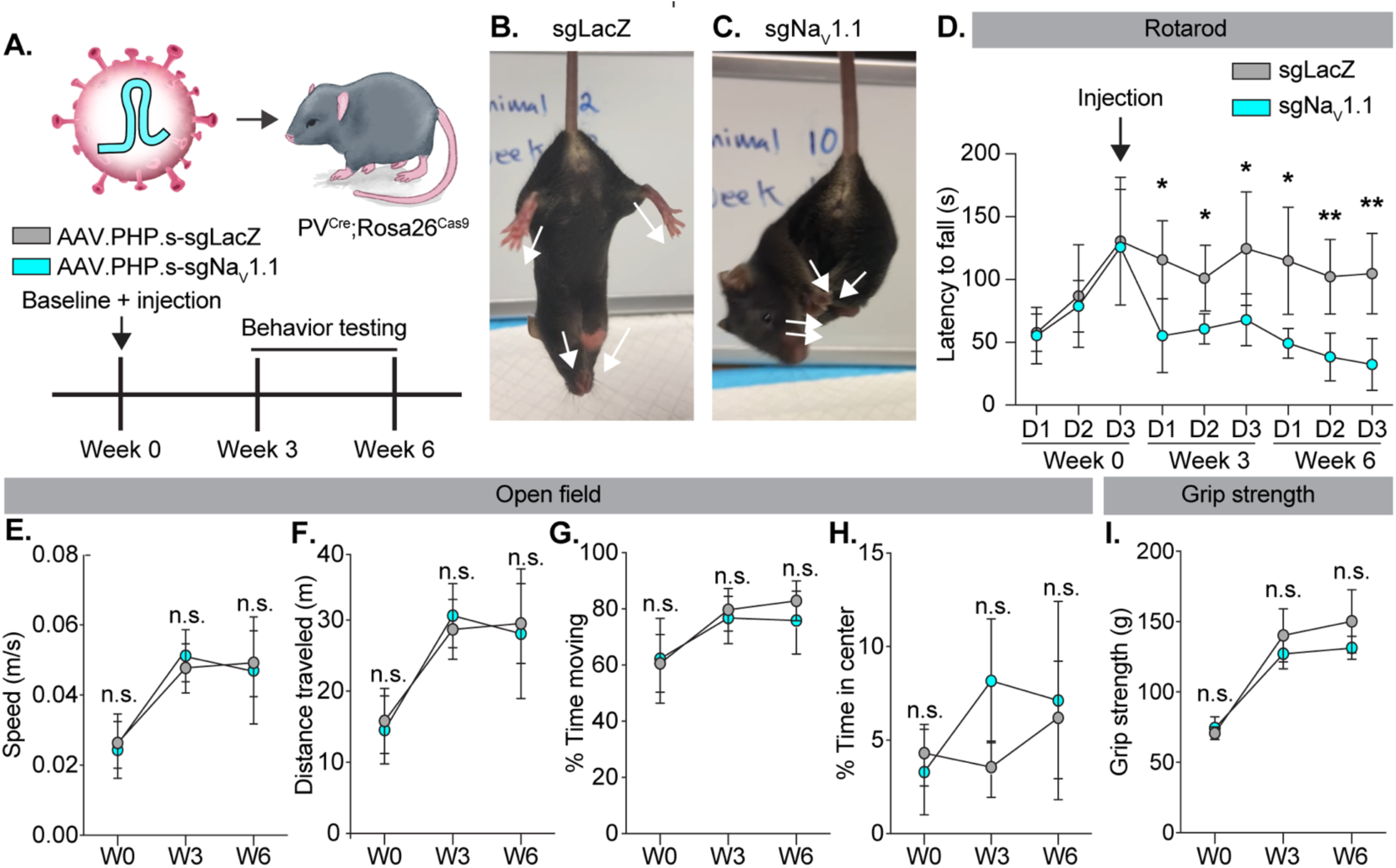
Na_V_1.1 is acutely required in proprioceptors for motor coordination. (A)Top, AAV.PHP.S virus containing sgRNAs targeting Na_V_1.1 (cyan) or LacZ (magenta) were retro-orbitally delivered to PV^Cas9^ mice. sgNa_V_1.1 is depicted in virus. Bottom, experimental timeline for behavioral testing. Baseline motor behavior assays were conducted at Week 0. Immediately following virus was delivered and subsequent behavioral testing was repeated 3- and 6-weeks post injection. Representative images showing abnormal limb position in sgLacZ (B) and sgNa_V_1.1 (C) arrows indicate direction of limbs. (D) Quantification of latency to fall on the rotarod across weeks. (E to H) Quantification of the open field parameters during a 10-minute open-field trail. (I) Average grip force in grams measured across six consecutive trials. A 2-way mixed design ANOVA (Fisher’s post hoc comparison) was used to determine statistical significance in D to I. N=5 sgLacZ, N=5 sgNa_V_1.1 N = mice.

This phenotype persisted up to 6 weeks following viral injection (Extended data Fig. 4-1), suggesting that the acute loss of Na_V_1.1 is not compensated for by other Na_V_ subtypes during this timeframe. In addition to rotarod, we also assayed spontaneous locomotion during a 10-minute open field trial (Fig. 4E-H) and also tested grip strength (Fig. 4I). We did not, however, observe a significant difference in performance between groups in these tests. Taken together, these experiments uncover an important role for Na_V_1.1 in proprioceptors for motor coordination.

### Targeting *Scn1a* disrupts action potential waveform in genetically identified proprioceptors

Compared to other sensory neurons subtypes, proprioceptors exhibit narrow action potentials which support high frequency spiking (Zheng et al., 2019). The intrinsic biophysical features of Na_V_1.1, such as rapid recovery from fast inactivation, resurgent, and persistent currents likely contribute to this feature (Kalume et al., 2007; Patel et al., 2015; Griffith et al., 2019). Thus, acute targeting of *Scn1a* in proprioceptors is expected to significantly disrupt action potential waveform. To address this, we packaged sgNa_V_1.1 or sgLacZ into the AAV.PHP.S-FLEX-TdTomato construct. This approach enabled genetic identification of PV-positive proprioceptors via TdT expression, facilitating whole-cell patch-clamp recordings *in vitro* when delivered to PV^Cas9^ mice 3 weeks after retro-orbital injection. We compared action potential waveforms from TdT-positive sensory neurons between sgNa_V_1.1 and sgLacZ groups. Single action potentials were elicited a depolarizing current injections in +50 pA increments to determine the rheobase current, defined as the minimal stimulus required to elicit an action potential (Fig. 5A). Phase plane analysis of single action potentials revealed differences in waveform dynamics between groups (Fig. 5B). Specifically, mice injected with sgNa_V_1.1 displayed a significant broadening of the action potential and a reduction in the maximal rate of rise (max dV/dt), consistent with reduced Na_V_ channel function (Fig. 5C-D). In addition, we observed a significant depolarizing shift in the minimal rate of voltage change (min dV/dt), reflecting altered repolarization dynamics (Fig. 5E). Because this measure reflects activation voltage-gated potassium (K_V_) channels, these findings suggest that reduced Na_V_ channel activity leads to concomitant decrease of K_V_ channel recruitment. Other intrinsic excitability parameters, including rheobase, action potential amplitude, and threshold were not significantly different between groups (Fig. 5F–H). Together, these data demonstrate the necessity of Na_V_1.1 for maintaining normal action potential kinetics in proprioceptors in adulthood.

**Figure 5.**
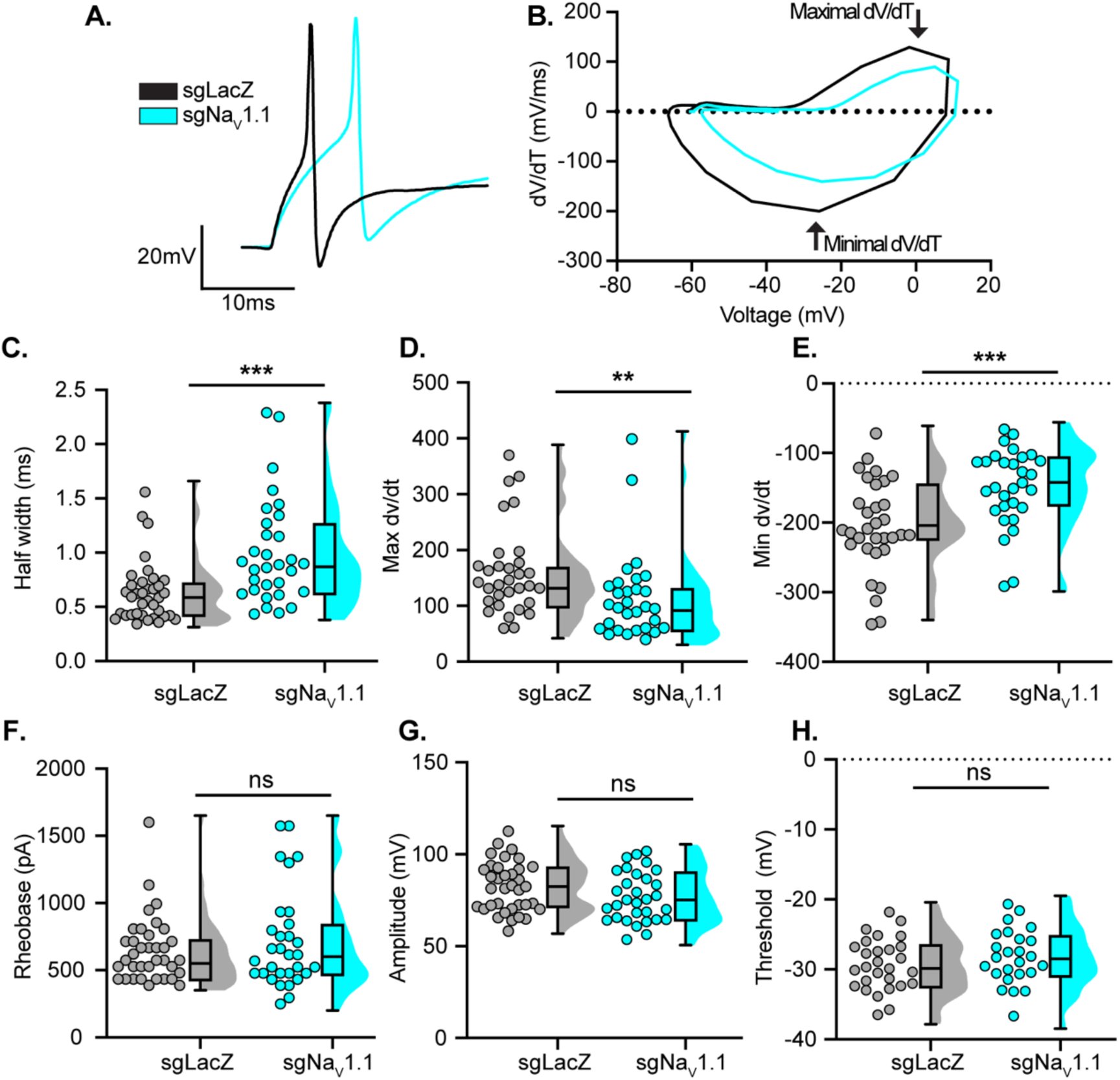
Acute targeting of *Scn1a* slows proprioceptor action potential kinetics. (A) Representative action potential waveforms from TdT-positive DRG neurons from sgLacZ (black) and sgNa_V_1.1 (cyan) injected mice. (B) Phase plane-plots of single action potentials shown in (A). Plots show the first derivative of the somatic membrane potential a function of somatic membrane potential. Quantification of the action potential half width (C), maximal and minimal rate of membrane potential change over time (D and E), Rheobase (F), action potential amplitude (G), and threshold (H). Significance was determined using a Mann-Whitney tests. Each dot represents a cell n=36 sgLacZ and n=31 sgNa_V_1.1 in C to G and n=28 sgLacZ and n=25 sgNa_V_1.1 in H. Data was collected from a total 6 biological replicates from each condition.

## Discussion

The development of transgenic tools and advances in single cell RNA sequencing have dramatically advanced our ability to genetically access specific cell types. These approaches have been integral to the investigation of ion channels and other proteins with cell-type-specific precision. Several caveats to this approach, however, remain. First, many neuronal populations share genetic markers and overlapping gene expression can complicate interpretation of behavioral measures when Cre recombinase is active in multiple cell types. This is especially true if the gene of interest is expressed across cell types. Second, in the absence of an inducible driver, Cre-mediated recombination is typically dependent on the timing of promoter activation, which limits temporal control and prevents insight into whether a gene is required during development or in adulthood. Lastly, while the use of triple transgenic lines that combine Cre and FlpO expression can circumvent some of these barriers, the generation and maintenance of such transgenic mouse lines imposes significant time and financial costs for many academic laboratories. Proprioceptors are one such example of technical limitations associated with traditional transgenic approaches. The genetic marker for proprioceptors in sensory neurons, PV, is also expressed broadly in other neuronal cell types notable for their role in motor function, like the cerebellum and spinal cord (Brandenburg et al., 2021; Veshchitskii and Merkulyeva, 2023). Furthermore, PV-expressing neurons across nervous systems, including those in the cerebellum and spinal cord, tend to rely on similar voltage-gated ion channel subtypes; (i.e., Na_V_1.1 and Na_V_1.6, Levin et al., 2006; Drouillas et al., 2023), in addition to also regulating motor function. Thus, traditional genetic approaches make it impossible to draw conclusions about the contributions of different ion channels to proprioceptor function at the behavioral level. These limitations highlight the need for alternative targeting strategies. To this end, we addressed this gap by developing an intersectional CRISPR/Cas9 gene editing approach that combines sensory-neuron–selective AAV capsid delivery of sgRNAs with PV-driven Cas9 expression. We successfully used this approach to disrupt Na_V_1.1 function *in vivo,* selectively in proprioceptors, with temporal control. We uncovered that Na_V_1.1 is acutely required in proprioceptors for motor behaviors and makes important contributions to the action potential waveform. Additionally, this approach represents a tractable platform for other laboratories investigating the function of specific genes in difficult to target sensory neuron populations.

Prior work had used the Cre/lox system to delete *Scn1a* in all peripheral sensory neurons (Pirt^Cre^;Na_V_1.1^fl/fl^, Na_V_1.1^cKO^), and found Na_V_1.1 expressed in these neurons to be essential for motor behaviors (Espino et al., 2022). While this work demonstrated a critical role for Na_V_1.1 in maintaining proprioceptor afferent excitability during static muscle stretch, a direct role for this channel in proprioceptor-driven behaviors could not be deduced due to non-selective targeting strategies. Indeed, Na_V_1.1 is also expressed in other sensory neuron populations involved in motor function, most notably low threshold mechanoreceptors (Gradwell et al., 2024). Using this intersectional CRISPR/Cas9 approach, we now demonstrate that disruption of *Scn1a* in adulthood leads to observable and quantifiable motor coordination deficits as early as 3 weeks post injection of sgNa_V_1.1 (Fig. 4D). These results now clarify the acute necessity of Na_V_1.1 in proprioceptors for motor behaviors. Interestingly, gross motor behaviors analyzed in the open-field and grip strength were unaffected by this manipulation (Fig. 4E-I). Conversely, prior work observed Na_V_1.1^cKO^ mice moved more slowly in the open field compared to floxed controls, despite spending the same amount of time moving (Espino et al., 2022). There are at least three interpretations for this result. First, based on viral transduction data, our intersectional model does not lead to Na_V_1.1 loss-of-function in all proprioceptors; therefore, the behavioral phenotypes observed here are less severe in comparison to Na_V_1.1^cKO^ mice. Alternatively, it is possible these differences arise due to Na_V_1.1 function in touch receptors for motor control. Indeed, prior work has demonstrated that cutaneous feedback modulates interneuron populations in the spinal cord for motor control (Gradwell et al., 2024), and Na_V_1.1 is widely expressed in cutaneous touch receptors (Zheng et al., 2019). Acute targeting of Na_V_1.1 selectively in touch receptors could help clarify the relative contributions of this channel to motor behaviors in different mechanoreceptor subtypes. Finally, it is likely that developmental loss of Na_V_1.1 in proprioceptors produces more severe phenotypes than this acute targeting strategy.

In our whole-cell patch-clamp recordings, we found that Na_V_1.1 loss-of-function led to a significant broadening of the action potential waveform in proprioceptors (Fig. 5). These findings are consistent with previous work using pharmacological blockers of Na_V_1.1 (Espino et al., 2022), as well as other studies investigating the role of Na_V_1.1 in fast-spiking neuronal populations (Ogiwara et al., 2007; Han et al., 2012). While these results support an acute role for Na_V_1.1 in proprioceptor electrogenesis, they also highlight the utility of this intersectional strategy as a new method for investigating Na_V_ channel function. Due to the high degree of sequence homology between Na_V_ channel isoforms, developing subtype-selective pharmacological agents remains a significant challenge. Additionally, constitutive genetic approaches are often confounded by developmental compensation, as the loss of one Na_V_ subtype can trigger upregulation of others (Ye et al., 2018; Bennett et al., 2019). The observation that sgNa_V_1.1 mice did not recover motor function, with their deficits persisting up to 6 weeks post injection suggests minimal, if any, compensation by other Na_V_ channel subtypes (Fig. 4C). Therefore, this intersectional approach may also serve as a useful tool for interrogating the function of specific ion channels in native cells with minimal compensatory confounds.

A notable limitation to our study is the analysis of indel frequency induced by sgNa_V_1.1 *in vivo*. We used FACS to isolate virally transduced TdT+ proprioceptors and found an indel percentage of only 7.95% and 6.8% in PV^Cas9^ and Pirt^Cas9^ mice, respectively. This observed indel rate is much lower than previous reports using CRISPR/Cas9 to target genes in central neurons, where editing efficiencies of up to 90% are reported (Hunker et al., 2020; Juarez et al., 2023). Given the robustness of our behavioral data, however, the most likely explanation for the reduced detection of indels *in vivo* is due to the heterogeneity of cell types present in DRG cell suspensions. Indeed, the presence of non-neuronal cells in our samples, which could be up to 50% based on prior reports (Tonello et al., 2023), could lead to an overrepresentation of wild type amplicons, and thus an underestimation of indel frequency induced by sgNa_V_1.1. Approaches using a nuclear FACS strategy to validate mutagenesis *in vivo* could address this technical limitation (Swiech et al., 2015; Hunker and Zweifel, 2020). Nevertheless, our *in vitro* screen (Fig. 1), selectivity of viral transduction (Fig. 2), observed motor coordination deficits (Fig. 4), and altered action potential kinetics (Fig. 5), together support a more robust rate of mutagenesis in the *Scn1a* gene in proprioceptors than detected in this study.

Collectively, we conclude that the use of this intersectional CRISPR/Cas9 strategy serves as reliable means for investigating the role of different genes of interest in somatosensory neurons.

## Acknowledgments

We would like to acknowledge other members of the Griffith Lab and the lab of Dr. Jorge Contreras for helpful feedback. We would also like to thank Josh Tulman for assistance with graphics. A subset of viral vectors used in this study were generated by the University of Minnesota Viral Vector and Cloning Core (Minneapolis, MN). Claude Opus 4.7 was used for language-editing during manuscript preparation.

## Extended data

**Extended data 1-1.**
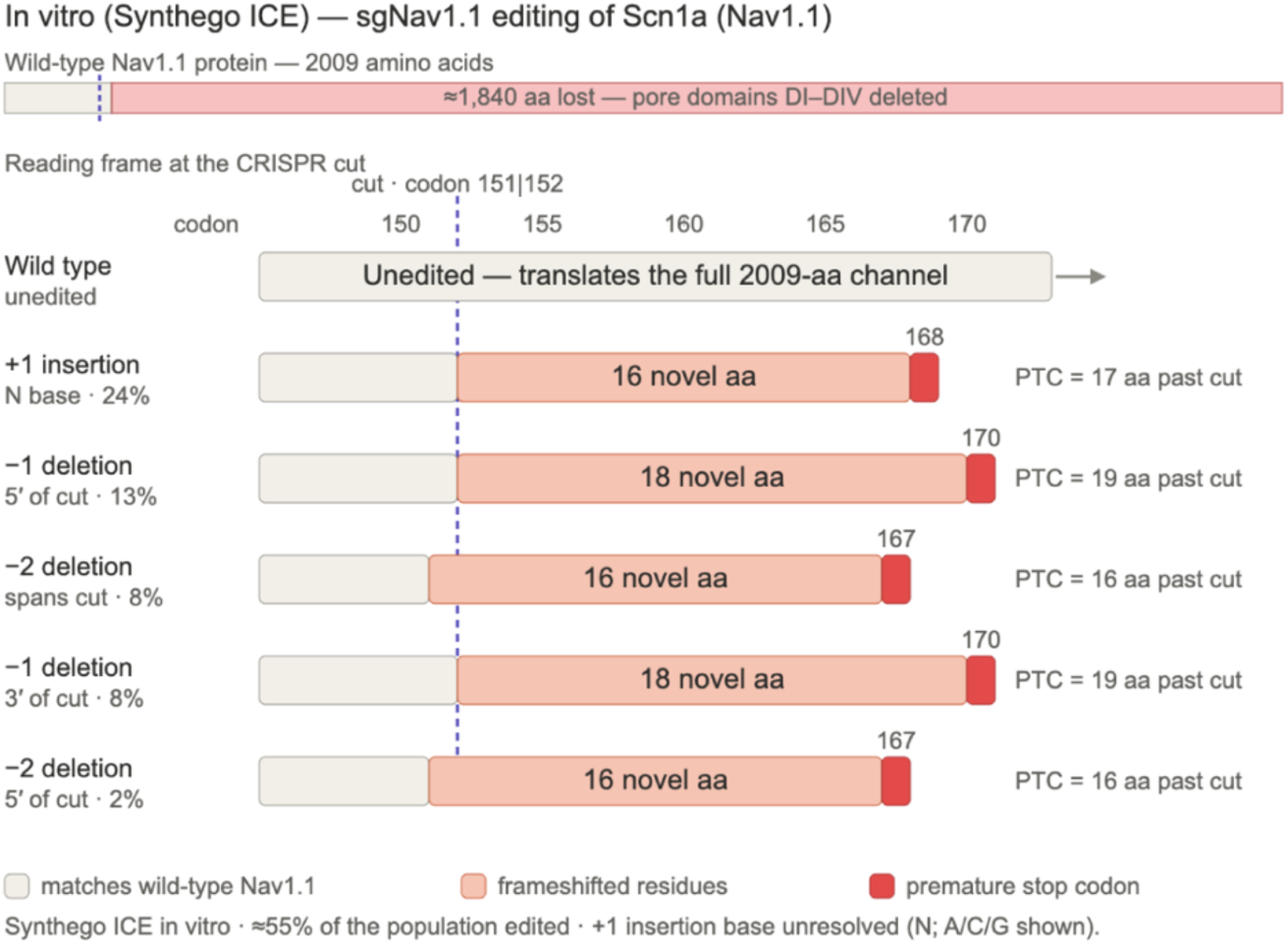
*In vitro* CRISPR editing of *Scn1a* with sgNa_V_1.1 generates frameshift alleles that truncate Nav1.1. Synthego ICE data of the sgNa_V_1.1-edited population *in vitro*, with each edited allele mapped onto the Na_V_1.1 coding sequence (*Scn1a*, NM_001313997.1; 2009-residue protein). The single guide directs a blunt SpCas9 cut at the codon 151/152 boundary (dashed line). Five edited alleles were resolved, shown with their relative contributions: a +1 insertion (24%), a −1 deletion 5′ of the cut (13%), a −2 deletion spanning the cut (8%), a −1 deletion 3′ of the cut (8%), and a −2 deletion 5′ of the cut (2%); ≈55% of the population carried an edit. Every allele is a frameshift, until it reaches a premature stop codon (PTC) 16–19 codons downstream of the cut (codon 167, 168, or 170). Segments matching wild-type Nav1.1 are shown in grey, frameshifted residues in salmon, and the premature stop codon in red. This figure was prepared with the assistance of Claude Cowork.

**Extended data 3-1.**
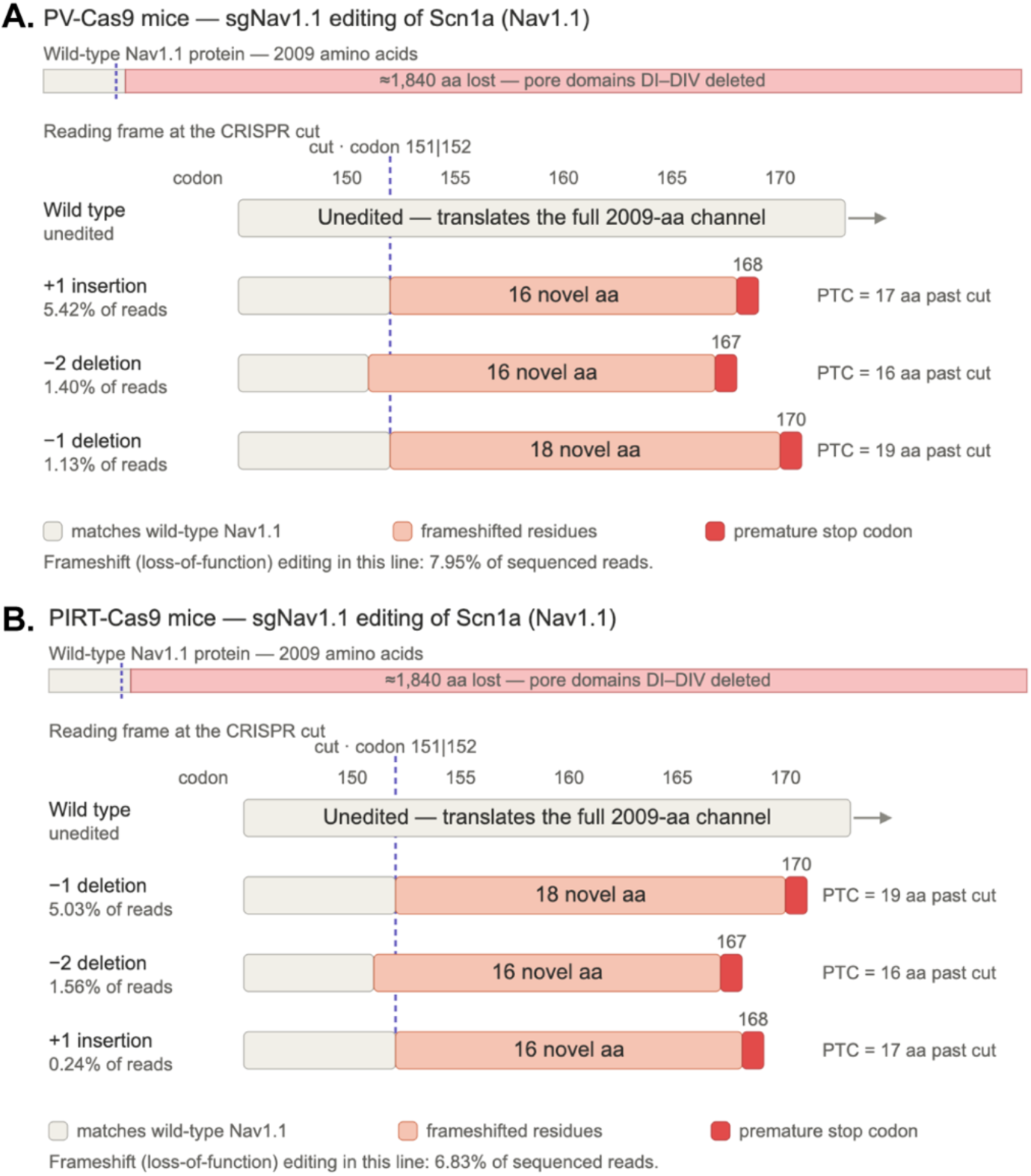
***In vivo* CRISPR editing of *Scn1a* with sgNa_V_1.1 generates frameshift alleles that truncate Na_V_1.1 in PV^Cas9^ and Pirt^Cas9^ mice.** Amplicon deep-sequencing of the sgNa_V_1.1-edited *Scn1a* locus in PV^Cas9^ (A) and Pirt^Cas9^ (B) with each edited allele mapped onto the Nav1.1 coding sequence (Scn1a, NM_001313997.1; 2009-residue protein). Three frameshift alleles were detected in each line. A +1 insertion, a −1 deletion, and a −2 deletion listed with their read frequencies and ordered by frequency. Every allele is a frameshift until it reaches a premature stop codon (PTC) 16–19 codons downstream of the cut. Segments matching wild-type Na_V_1.1 are shown in grey, frameshifted residues in salmon, and the premature stop codon in red. Total frameshift (loss-of-function) editing was 7.95% of reads in PV^Cas9^ and 6.83% in Pirt^Cas9^. This figure was prepared with the assistance of Claude Cowork.

**Extended data 4-1.**
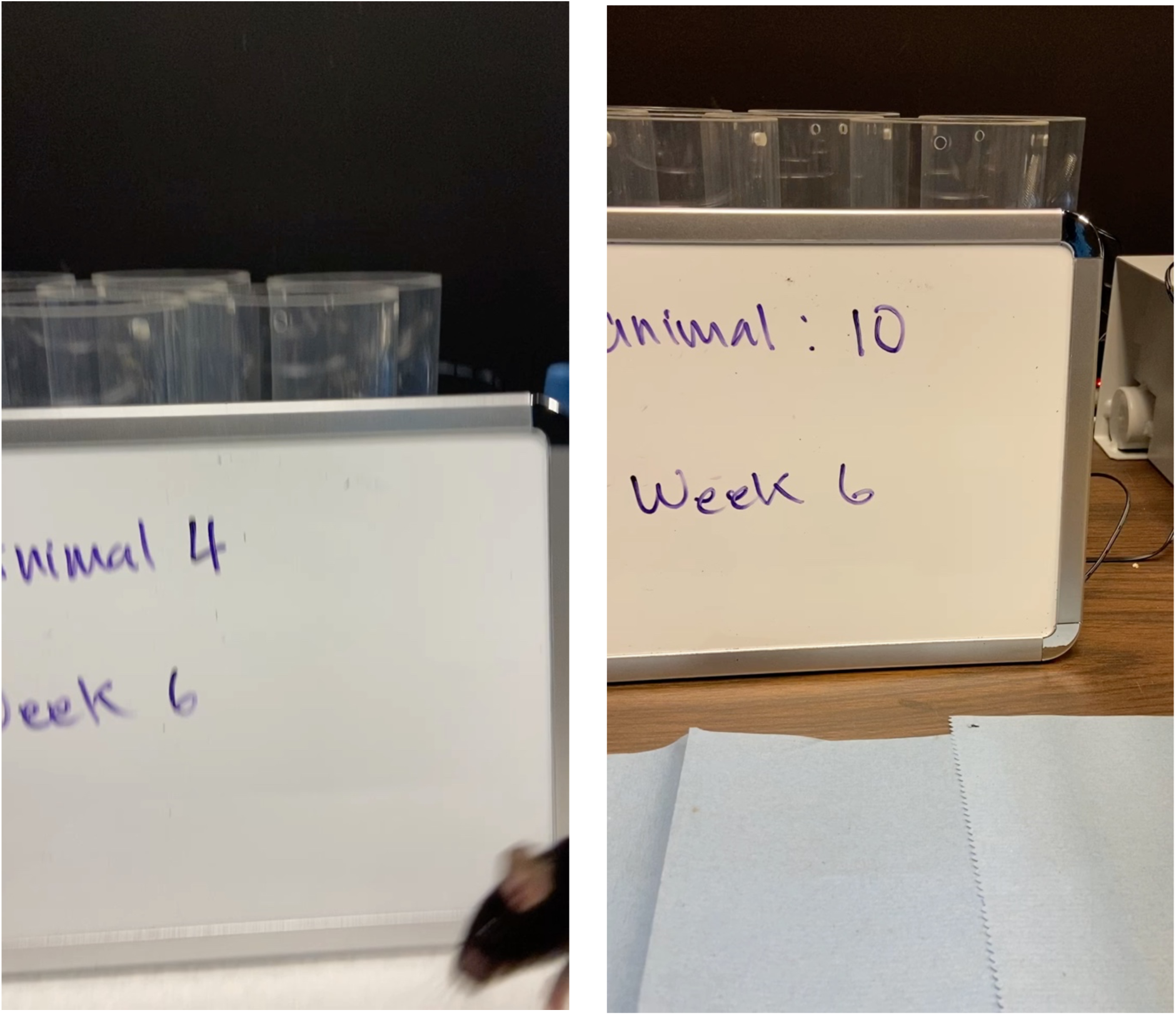
Acute targeting of *Scn1a* in proprioceptors leads to visible behavioral impairments. Representative motor hind limb phenotype between a PVCas9 mice injected with either control sgLacZ (Left) or sgNa_V_1.1 (Right)

## REFERENCES

Asencor AI, Dvoryanchikov G, Makhoul V, Tsoulfas P, Chaudhari N (2022) Selectively Imaging Cranial Sensory Ganglion Neurons Using AAV-PHP.S. eNeuro 9 Available at: 10.1523/ENEURO.0373-21.2022.

Bennett DL, Clark AJ, Huang J, Waxman SG, Dib-Hajj SD (2019) The Role of Voltage-Gated Sodium Channels in Pain Signaling. Physiol Rev 99:1079–1151.

Brandenburg C, Smith LA, Kilander MBC, Bridi MS, Lin Y-C, Huang S, Blatt GJ (2021) Parvalbumin subtypes of cerebellar Purkinje cells contribute to differential intrinsic firing properties. Mol Cell Neurosci 115:103650.

Chan KY, Jang MJ, Yoo BB, Greenbaum A, Ravi N, Wu W-L, Sánchez-Guardado L, Lois C, Mazmanian SK, Deverman BE, Gradinaru V (2017) Engineered AAVs for efficient noninvasive gene delivery to the central and peripheral nervous systems. Nat Neurosci 20:1172–1179.

Drouillas B, Brocard C, Zanella S, Bos R, Brocard F (2023) Persistent Nav1.1 and Nav1.6 currents drive spinal locomotor functions through nonlinear dynamics. Cell Rep 42:113085.

Espino CM, Lewis CM, Ortiz S, Dalal MS, Garlapalli S, Wells KM, O’Neil DA, Wilkinson KA, Griffith TN (2022) NaV1.1 is essential for proprioceptive signaling and motor behaviors. Elife 11 Available at: 10.7554/eLife.79917.

Ferguson B, Glick C, Huguenard JR (2023) Prefrontal PV interneurons facilitate attention and are linked to attentional dysfunction in a mouse model of absence epilepsy. Elife 12 Available at: 10.7554/eLife.78349.

Gradwell MA et al. (2024) Multimodal sensory control of motor performance by glycinergic interneurons of the mouse spinal cord deep dorsal horn. Neuron 112:1302–1327.e13.

Griffith TN, Docter TA, Lumpkin EA (2019) Tetrodotoxin-Sensitive Sodium Channels Mediate Action Potential Firing and Excitability in Menthol-Sensitive Vglut3-Lineage Sensory Neurons. J Neurosci 39:7086–7101.

Han S, Tai C, Westenbroek RE, Yu FH, Cheah CS, Potter GB, Rubenstein JL, Scheuer T, de la Iglesia HO, Catterall WA (2012) Autistic-like behaviour in Scn1a+/- mice and rescue by enhanced GABA-mediated neurotransmission. Nature 489:385–390.

Hunker AC, Soden ME, Krayushkina D, Heymann G, Awatramani R, Zweifel LS (2020) Conditional Single Vector CRISPR/SaCas9 Viruses for Efficient Mutagenesis in the Adult Mouse Nervous System. Cell Rep 30:4303–4316.e6.

Hunker AC, Zweifel LS (2020) Protocol to design, clone, and validate sgRNAs for In Vivo reverse genetic studies. STAR Protoc 1:100070.

Juarez B, Kong M-S, Jo YS, Elum JE, Yee JX, Ng-Evans S, Cline M, Hunker AC, Quinlan MA, Baird MA, Elerding AJ, Johnson M, Ban D, Mendez A, Goodwin NL, Soden ME, Zweifel LS (2023) Temporal scaling of dopamine neuron firing and dopamine release by distinct ion channels shape behavior. Sci Adv 9:eadg8869.

Kalume F, Yu FH, Westenbroek RE, Scheuer T, Catterall WA (2007) Reduced sodium current in Purkinje neurons from Nav1.1 mutant mice: implications for ataxia in severe myoclonic epilepsy in infancy. J Neurosci 27:11065–11074.

Khlghatyan J, Beaulieu J-M (2020) CRISPR-Cas9-Mediated Intersectional Knockout of Glycogen Synthase Kinase 3β in D2 Receptor-Expressing Medial Prefrontal Cortex Neurons Reveals Contributions to Emotional Regulation. CRISPR J 3:198–210.

Kim AY, Tang Z, Liu Q, Patel KN, Maag D, Geng Y, Dong X (2008) Pirt, a phosphoinositide-binding protein, functions as a regulatory subunit of TRPV1. Cell 133:475–485.

Lehnert BP, Santiago C, Huey EL, Emanuel AJ, Renauld S, Africawala N, Alkislar I, Zheng Y, Bai L, Koutsioumpa C, Hong JT, Magee AR, Harvey CD, Ginty DD (2021) Mechanoreceptor synapses in the brainstem shape the central representation of touch. Cell 184:5608–5621.e18.

Levin SI, Khaliq ZM, Aman TK, Grieco TM, Kearney JA, Raman IM, Meisler MH (2006) Impaired motor function in mice with cell-specific knockout of sodium channel Scn8a (NaV1.6) in cerebellar purkinje neurons and granule cells. J Neurophysiol 96:785–793.

Maizels N (2013) Genome engineering with cre-loxP. J Immunol 191:5–6.

McLellan MA, Rosenthal NA, Pinto AR (2017) Cre-loxP-mediated recombination: General principles and experimental considerations: Cre-loxP-mediated recombination. Curr Protoc Mouse Biol 7:1–12.

Meisler MH, O’Brien JE, Sharkey LM (2010) Sodium channel gene family: epilepsy mutations, gene interactions and modifier effects: Sodium channel gene family. J Physiol 588:1841–1848.

Ogiwara I, Miyamoto H, Morita N, Atapour N, Mazaki E, Inoue I, Takeuchi T, Itohara S, Yanagawa Y, Obata K, Furuichi T, Hensch TK, Yamakawa K (2007) Nav1.1 localizes to axons of parvalbumin-positive inhibitory interneurons: a circuit basis for epileptic seizures in mice carrying an Scn1a gene mutation. J Neurosci 27:5903–5914.

Park Y-N, Masison D, Eisenberg E, Greene LE (2011) Application of the FLP/FRT system for conditional gene deletion in yeast Saccharomyces cerevisiae. Yeast 28:673–681.

Patel RR, Barbosa C, Xiao Y, Cummins TR (2015) Human Nav1.6 Channels Generate Larger Resurgent Currents than Human Nav1.1 Channels, but the Navβ4 Peptide Does Not Protect Either Isoform from Use-Dependent Reduction. PLoS One 10:e0133485.

Proske U, Gandevia SC (2012) The proprioceptive senses: their roles in signaling body shape, body position and movement, and muscle force. Physiol Rev 92:1651–1697.

Ran FA, Hsu PD, Wright J, Agarwala V, Scott DA, Zhang F (2013) Genome engineering using the CRISPR-Cas9 system. Nat Protoc 8:2281–2308.

Sharma N, Flaherty K, Lezgiyeva K, Wagner DE, Klein AM, Ginty DD (2020) The emergence of transcriptional identity in somatosensory neurons. Nature 577:392–398.

Swiech L, Heidenreich M, Banerjee A, Habib N, Li Y, Trombetta J, Sur M, Zhang F (2015) In vivo interrogation of gene function in the mammalian brain using CRISPR-Cas9. Nat Biotechnol 33:102–106.

Tonello R, Silveira Prudente A, Hoon Lee S, Faith Cohen C, Xie W, Paranjpe A, Roh J, Park C-K, Chung G, Strong JA, Zhang J-M, Berta T (2023) Single-cell analysis of dorsal root ganglia reveals metalloproteinase signaling in satellite glial cells and pain. Brain Behav Immun 113:401–414.

Veshchitskii A, Merkulyeva N (2023) Calcium-binding protein parvalbumin in the spinal cord and dorsal root ganglia. Neurochem Int 171:105634.

Wang D, Zhang F, Gao G (2020) CRISPR-based therapeutic genome editing: Strategies and in vivo delivery by AAV vectors. Cell 181:136–150.

Wilkinson KA (2022) Molecular determinants of mechanosensation in the muscle spindle. Curr Opin Neurobiol 74:102542.

Wu S-X, Koshimizu Y, Feng Y-P, Okamoto K, Fujiyama F, Hioki H, Li Y-Q, Kaneko T, Mizuno N (2004) Vesicular glutamate transporter immunoreactivity in the central and peripheral endings of muscle-spindle afferents. Brain Res 1011:247–251.

Yarmolinsky M, Hoess R (2015) The legacy of Nat Sternberg: The genesis of Cre-lox technology. Annu Rev Virol 2:25–40.

Ye M, Yang J, Tian C, Zhu Q, Yin L, Jiang S, Yang M, Shu Y (2018) Differential roles of NaV1.2 and NaV1.6 in regulating neuronal excitability at febrile temperature and distinct contributions to febrile seizures. Sci Rep 8:753.

Zhang X, Yu X, Tuo M, Zhao Z, Wang J, Jiang T, Zhang M, Wang Y, Sun Y (2023) Parvalbumin neurons in the anterior nucleus of thalamus control absence seizures. Epilepsia Open 8:1002–1012.

Zheng Y, Liu P, Bai L, Trimmer JS, Bean BP, Ginty DD (2019) Deep Sequencing of Somatosensory Neurons Reveals Molecular Determinants of Intrinsic Physiological Properties. Neuron 103:598–616.e7.

